# Biopriming of broad bean seeds with entomopathogenic fungus *Metarhizium robertsii* does not affect invertebrate communities of the agroecosystem

**DOI:** 10.1101/2022.07.20.500881

**Authors:** I.I. Lyubechanskii, T.A. Novgorodova, Y. Panina, V.V. Kryukov, V.S. Sorokina, T.A. Sadokhina, D.Ju. Bakshaev, R.Ju. Dudko, A.A. Gurina, V.V. Glupov

## Abstract

Biopriming, or treatment of seeds with beneficial microorganisms such as beneficial fungi, can be a promising strategy in agricultural cultivation. However, the effects of such treatment on non-target organisms living in the soil and on plants have not been sufficiently studied, and it is not known very well whether such treatment would alter invertebrate communities (e.g., harm them). Here, we addressed the effect of treating broad bean seeds (*Vicia faba* L.) with the conidia of entomopathogenic ascomycete *Metarhizium robertsii* on the diversity and abundance of invertebrate communities in the agroecosystem in the south part of West Siberia in 2019 and 2020. We have analyzed the effect both on the general invertebrate communities as well as on the main pests of beans. In the case of bean pests, we assessed the rate of plant infestation by aphids (Hemiptera: Aphididae) and the degree of leaf damage by leafminer flies *Liriomyza bryoniae* (Diptera: Agromyzidae). In most cases, the treatment did not lead to significant changes in the total abundance of the soil invertebrates and herbivores or the abundance of predominant taxa (Coleoptera: Carabidae, Staphylinidae, Elateridae, Scarabaeidae, Curculionidae; Hemiptera: Miridae, Cicadellidae, Aphididae; larvae of Diptera). A positive effect of treatment on population density of the soil mesofauna was noted for Diptera larvae in June 2019. Regarding aphids and leafminer flies, no significant effect was observed in terms of the proportion of plants with aphids and the density of aphid colonies on individual plants throughout the season, and no significant influence was found on the proportion of plant leaves damaged by leafminer fly *Liriomyza bryoniae* larvae. In summary, in Western Siberia, the treatment of broad bean seeds with *M. robertsii* did not significantly affect non-target arthropods common for bean fields as well as the main pests of beans, namely aphids and miner flies.

## Introduction

Entomopathogenic ascomycetes *Metarhizium* are common soil fungi. They develop on host insects and interact with different plants as unspecialized rhizosphere colonizers and endophytes (1,2). It is a well-known fact that biopriming using fungi may lead to many desirable effects on plants, such as the promotion of growth, nitrogen and phosphorus delivery, modulation of immunity, increased defense against phytopathogens and pest insects (3–5). Moreover, plant colonization by the entomopathogenic fungi may lead to changes in the abundance of herbivore arthropods through a variety of factors, such as direct contacts with fungal propagules, an influence of fungal toxins on arthropods when they are feeding on fungi-colonized plants, biochemical changes in colonized plants, influence on plant biomass and terms of their development etc. Negative effects on arthropods are mainly manifested as disturbances in arthropods’ feeding, decrease in larval weight and survival, drop in fecundity, changes in host choice and decrease in abundance (6). It is well known that treatment of plants with entomopathogenic fungi (for example by inoculation of seeds) leads to their subsequent dispersal in the root system and rhizosphere soil (7). Therefore, an increase in abundance of fungi may influence non-target arthropods in the soil.

Previous studies on the treatment of plants with entomopathogenic fungi have shown inconsistent effects on phytophages (1,8,9). This can be explained by differences in environmental conditions and experimental methods in these studies. For example, investigations in systems “entomopathogenic fungi – plants – insects” were mostly conducted in laboratory and greenhouse conditions, with only a few field experiments (1). Inconsistent effects may also be caused by differences in trophic relationship between arthropods, plants and endophytic fungi. The fungal community in plants and in the soil of the rhizosphere is always represented by a wide range of non-entomopathogenic fungi, mainly phytopathogens and saprotrophs (10–12). In a recent meta-analysis, Gange and coworkers (6) showed that entomopathogenic (and non-entomopathogenic) endophytes have negative effects on insects from various orders, with the most pronounced effects on sucking insects Hemiptera, Thysanoptera and Hymenoptera. Interestingly, entomopathogenic endophytes negatively affect insects with different breadth of trophic preferences (polyphages, oligophages, and monophages), while non-entomopathogenic endophytes affect polyphages and oligophages, but do not affect monophages. It should also be noted that the effect of entomopathogenic fungi on non-target arthropods was studied mainly under the conditions of direct contacts of the invertebrates with fungal propagules (direct inoculation or inoculation of the environment with fungi) as well as of feeding with fungal metabolites (13–16). Little is known about the effects on rhizosphere or endophytic colonization of plants by entomopathogenic fungi on non-herbivore invertebrates.

Regarding the colonization of broad beans (*Vicia faba* L.) with entomopathogenic / endophytic fungi and its influence on insects, several assays have been performed in laboratory conditions using sterile substrates. Specifically, Jaber and Enkerli (17) registered successful systemic colonization of *V. faba* after soaking the seeds in the suspension with both *Beauveria bassiana* and *M. brunneum* and subsequently growing them on sterile substrates, however, the effect on arthropods was not assessed. Akello and Sikora (18) investigated endophytic colonization of *V. faba* with *B. bassiana* and *M. anisopliae* on sterile soil and showed negative effects on aphids *Acyrthosiphon pisum* and *Aphis fabae*. Akutse and coworkers (19) used sterile manure mixture and demonstrated a negative effect of *V. faba* colonization with *B. bassiana* on leafminer *Liriomyza huidobrensis* (Diptera) and its parasitoids *Phaedrotoma scabriventris* and *Diglypus isaea* (Hymenoptera). However, to our knowledge, there are no studies on inoculation of *V. faba* with entomopathogenic fungi that would assess its effects on target and non-target invertebrates in field conditions. In addition, even though many studies have focused on assessing the effects found after treatment of plants with entomopathogenic fungi (20,21), the effects of seed treatment with entomopathogenic fungi on the structure of invertebrate communities are still insufficiently studied, and the degree of possible negative impact on non-target invertebrates may be underestimated.

This work is a part of a comprehensive study of the effect of endophytes on the entire agricultural system of the broad bean field, including agricultural plants, fungi and invertebrates, both pests and accompanying species. The characteristics of the colonization of plant tissues and rhizosphere soil by the fungus *M. robertsii*, as well as the effects on plants, are detailed in our previous work (22). Briefly, in 2019, the level of colonization on the plots treated with *M. robertsii* did not differ from the control. In 2020, fungi treatment resulted in a 2-fold increase in the number of CFU (colony-forming units) of *M. robertsii* in the rhizosphere soil in June-July (p<0,05), followed by a decline in August to the control level. In June, the colonization of internal tissues of stems and root surfaces reached 18% and 36%, respectively, compared to 4% of *Metarhizium*-positive plants in the control. In the following months (July-August), the level of colonization decreased to the control values. It was found that soaking broad bean seeds with *M. robertsii* conidia before planting led to significant decrease in the intensity of plant diseases (root rot, powdery mildew, spotting) even at a relatively low level of plant colonization by the fungus. In addition, this treatment stimulated the formation of active nodules on the roots, increased plant biomass and the yield of beans in agroecosystems in Western Siberia.

The aim of this work was to investigate the effect of treatment of broad bean seeds with *M. robertsii* conidia on the composition and structure of invertebrate communities in the agroecosystem. The goals of this study were as follows: (1) to assess the impact of processing on the soil mesofauna (soil inhabitants and soil surface inhabitants), as well as insects – inhabitants of the grass stand (herbivores); (2) to assess the impact of treatment on the main pests of beans, namely, the level of plant infestation by aphids, and the extent of leaf damage by miner flies.

## Materials and methods

### Plant and fungal material

Broad beans (*Vicia faba* L.) of breed “Sibirskii” were chosen as a model plant culture. *Metarhizium robertsii* (strain P-72, Genbank #KP172147) from the collection of microorganisms of the Institute of Systematics and Ecology of Animals Siberian Branch of the Russian Academy of Sciences (ISEA SB RAS) was used in the study. Fungal conidia were cultivated on autoclaved millet as previously described (23). Concentrations of conidia were determined using a Neubauer hemocytometer. The viability of conidia was checked by plating of suspensions on Sabouraud dextrose agar. Germination percentage was 95%.

### Location and experimental setup

Field trials were carried out over two summer seasons in 2019 and 2020 in the forest-steppe zone of Western Siberia in the vicinity of Novosibirsk (Krasnoobsk settlement; 54°55ʹ N, 82°59ʹ E) at the field station of the Siberian Federal Scientific Centre of Agro-BioTechnologies of the Russian Academy of Sciences (SFSCA RAS). The soil of the experimental site consists of leached chernozem, medium-thick, medium-loamy. Experimental plots (10×3.9 m^2^ for each plot) were established, where the effect of treating beans seeds with the entomopathogenic fungus *Metarhizium robertsii* on invertebrate communities was assessed in comparison with the control. In 2019, the experiment was carried out in four replicates (a total of 8 plots were established), and in 2020 the experiment was carried out in five replicates (a total of 10 plots).

### Seed treatment, planting and main effects on plants

The treatment of plants was carried out by soaking the seeds according to the previously described method (22). Conidia were suspended in a water-tween solution (Tween-20, 0.04%) with a final concentration of 5 × 10^7^ conidia/ml. The seeds of the broad beans were treated with the suspension (2.5 liters per 20 kg of the broad beans) and dried overnight before planting. The control was treated with a conidia-free water-tween solution. The planting of beans was carried out on May 16^th^ in 2019, and on May 19^th^ in 2020, when the soil temperature at a depth of 6 – 8 cm reached 8 – 10° C. The beans were planted using an Optima seeder (Kverneland Group Soest, GmbH) in one tier. The seeding depth was 6-8 cm, the width between the rows was 70 cm, with a seeding rate of 400 thousand germinable beans per hectare. Harvesting took place on October 10^th^ in 2019 and September 18^th^ in 2020.

### Collecting of invertebrates

#### Soil mesofauna and herbivores

To determine the homogeneity of the spatial distribution of the soil mesofauna (geo- and herpetobionts) on the experimental plots in May, a preliminary count of invertebrates was carried out immediately before planting seeds (2019) or before plowing the field (2020). In 2019 (May 14^th^), 10 soil samples (25×25×10 cm^3^) were taken from randomly distributed spots on the field, and in 2020 (May 12^th^), 20 samples (20×20×5 cm^3^) were collected according to a “checkerboard” pattern with distances between samples of about 4 m. To investigate the taxonomic composition and abundance of invertebrates, counting of invertebrates using a modified method of soil sampling (24,25) was carried out twice during the season (in 2019, on June 20^th^ and 25^th^ and on August 14^th^ and 16^th^; in 2020 on June 22^nd^ and 23^rd^, and on August 12^th^ and 13^th^). Soil samples with the size of 25×25×10 cm^3^ with bean plants (from 3 to 5, usually 4) on them were packed in ventilated bags made of synthetic fabric, and then the contents were disassembled by hand, while examining both the soil and the plants. This technique makes it possible to take into account the number of both soil inhabitants and the population of the aboveground parts of plants. In order to reduce the influence of the uneven distribution of invertebrates, three samples were taken from each plot in each of the counts. Thus, the total number of soil samples was 48 in 2019 and 60 in 2020.

The collected invertebrates were placed in 70% ethanol solution and then classified in the laboratory up to large taxa (orders in the case of annelids, arachnids, millipedes, insects other than Coleoptera; or families in the case of Coleoptera). Several sources were used to identify the material (26,27).

#### Aphids

Aphids were counted three times per season during different periods of plant vegetation: the branching phase (on June 11^th^, 2019, and on June 3^rd^, 2020; hereinafter referred to as “June”), the budding phase (on July 23^rd^, 2019, and on July 15^th^, 2020; hereinafter referred to as “July”) and the seed maturation phase (on August 30^th^, 2019, and on August 17^th^, 2020; hereinafter referred to as “August”). At each site, 10 randomly selected plants were examined. The number of aphids per plant and the proportion of plants infested with aphids were counted. Insects were collected into the 70% alcohol solution for further species identification. In total, during the study, 540 plants were examined (240 in 2019 and 300 in 2020), 1575 aphids were collected (217 in 2019 and 885 in 2020). Analysis of the material and further identification of aphids was carried out using Stemi 2000-C and Zeiss Axiostar Plus microscopes. Aphid slide mounts were made using the Faure-Berlese mounting medium. When identifying aphids, an Internet resource was used based on the works of Blackman and Eastop (28–30) which includes current information on aphids (31). Synonymy was given according to Favret (32). All materials were transferred and are currently stored at the Institute of Systematics and Ecology of Animals, Siberian Branch of the Russian Academy of Sciences (Novosibirsk, Russia).

#### Miner flies

To determine the level and dynamics of plant infestation with miner flies, the damaged foliage was counted twice per season in 2019 and three times in 2020, during different periods of plant vegetation: the branching phase (on June 3^rd^, 2020), the budding phase (on July 23^rd^, 2019, and on July 15^th^, 2020) and the seed maturation phase (on August 30^th^, 2019, and on August 17^th^, 2020). At each site, 10 randomly selected plants were examined. The leaf blades were examined for the presence of characteristic damage (mines). Both mines with larvae and those that were empty at the time of the survey were taken into account.

In order to assess the level of infestation of bean plants at different stages of development, the number of mines on each infected leaf was counted, and the proportion of infected leaves of a certain zone of the plant was assessed, specifically, up to the peduncles (hereinafter “the lower part of the plant”) in July 2019 and June 2020, the part of the plant with peduncles (hereinafter “the middle part”) in July 2019 and 2020, and above the peduncles (hereinafter “the upper part of the plant”) in August 2019 and 2020. Due to the impossibility of assessment in June 2019, the lower and middle parts of the plant were examined simultaneously during the investigation in July 2019. In total, 160 plants were examined in 2019, and 300 plants in 2020. To determine the species of miners, at each stage of plant examination, leaves with relatively fresh lesions characteristic for each species of miner flies were selected and placed in plastic containers for hatching adult flies (Fig. 1).

**Fig. 1.**
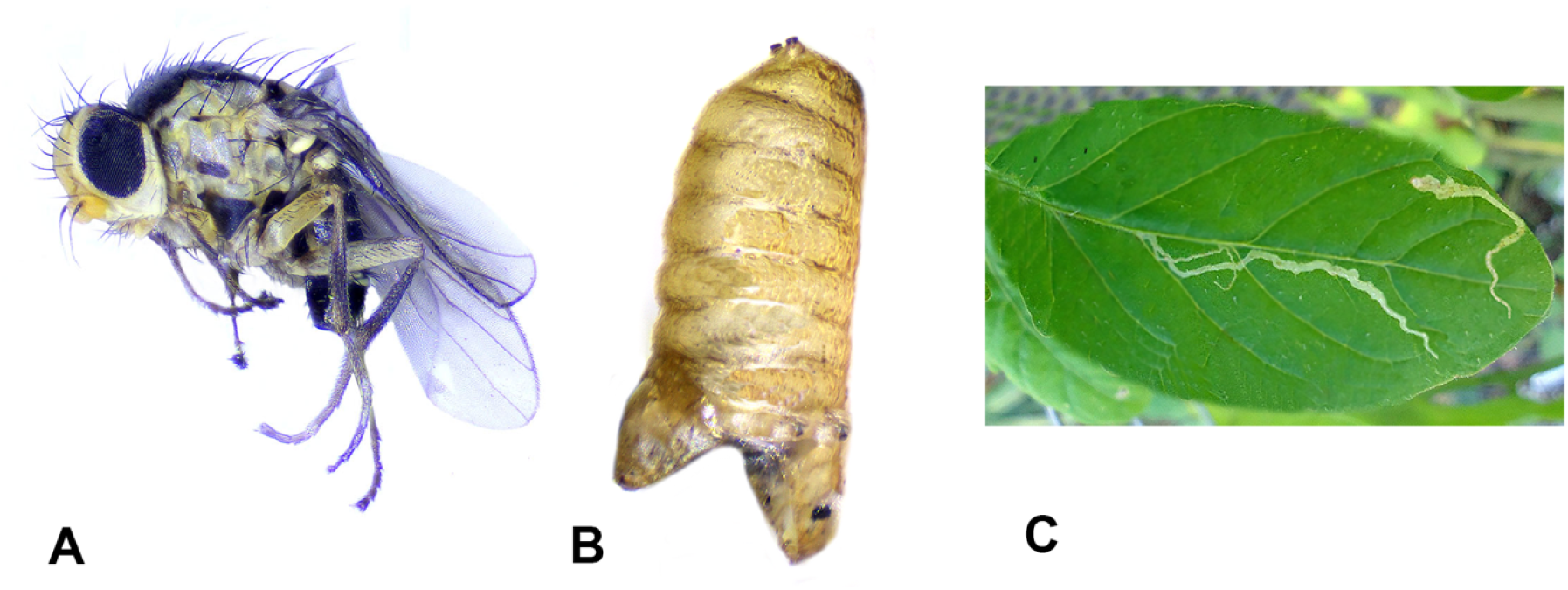
Leafminer *Liriomyza bryoniae*. A) Adult, lateral view. B) Open pupa; C) Mines on the leaves of beans.

The larvae were grown in the laboratory at room temperature (20-24°C), natural light and regular moisture (60-70%). In 2019, 24 larvae pupated, out of which 6 adult flies and 11 parasitoids developed, while 7 puparia died. In 2020, 23 larvae pupated, from which 6 adult flies and 10 parasitoids developed, while 7 puparia died. Identification of adult flies was carried out by examining external appearance using an Altami PSO745-T stereomicroscope. Preparations of male genitalia were made according to generally accepted techniques for Diptera.

#### Statistics

For the soil mesofauna and the complex of herbivores, the analysis of the influence of the type of treatment on the number of invertebrates was carried out using the examples of the most numerous taxa, from 7 to 9 at different times. The larvae and imagoes of Coleoptera and Neuroptera (lacewings) were counted and analyzed separately. Groups of invertebrates, in which the total number for all experimental sites did not exceed 10 specimens per count, were taken into account only in the comparative analysis of the number of taxa and the total number of invertebrates at the plots with different types of seed treatment. The variants of the invertebrate populations at each experimental site were ordered using multidimensional scaling after calculating the Euclidean distance. For the comparative analysis of the structure of invertebrate communities formed on plots with different treatments, the Shannon information index, the Simpson and Berger-Parker diversity indices (33), and the species evenness (34) were calculated. The significance of the difference was assessed using a permutation test (n = 9999).

The distribution of the studied parameters differed from normal (Shapiro-Wilk criterion, p<0.05), therefore, the Mann-Whitney test was used to estimate the effect of seed treatment on the composition (number of taxa) and abundance of soil mesofauna and herbivores (both in total and for individual groups), as well as on the number of aphids on plants, the proportion of plants infested by aphids, and the level of plant infection by miner fly larvae (proportion of leaves damaged by mines) at different phases of development. Analyses were performed using STATISTICA v.8.0.725, PAST 4 (35), and Microsoft Excel.

## Results

### Spatial heterogeneity of the population and population density of soil mesofauna and herbivores before and after treatment with *Metarhizium robertsii*

Before the start of the experiment (treatment with *Metarhizium robertsii*), representatives of 10 taxonomic groups of invertebrates were found in the soil samples in May 2019 and representatives of 9 taxonomic groups were found in May 2020 (Table 1).

**Table 1.** Population density of soil invertebrates at the experimental site before the start of the experiment.

| Total,<br>descriptive<br>statistics | May 2019 |  | May 2020 |  |
| --- | --- | --- | --- | --- |
|  | Specimens | taxa | Specimens | taxa |
| Min | 1 | 1 | 0 | 0 |
| Max | 22 | 6 | 6 | 4 |
| Median | 5 | 3 | 1 | 1 |
| 25 % | 2,5 | 1 | 0 | 0 |
| 75 % | 9,25 | 5,25 | 3 | 2 |
| Sum | 69 | 10 | 35 | 9 |
| Shapiro-Wilk W | 0,82 | 0,86 | 0,80 | 0,83 |
| p (normal) | 0,025 | 0,087 | 0,001 | 0,002 |
| **Groups of invertebrates** |  |  |  |  |
| Enchytraeidae | 2 [1; 3] |  | 2 [1; 2] |  |
| Araneae | 2 [1; 2] |  | 1 [1; 1,75] |  |
| Elateridae | 1 [1; 1,25] |  | 0 |  |
| Elateridae<br>(larvae) | 1,5 [1; 2] |  | 1 [1; 2] |  |
| Scarabaeidae | 1 [1; 1] |  | 0 |  |
| Carabidae | 1,5 [1; 2,75] |  | 1 [1; 1] |  |
| Carabidae<br>(larvae) | 0 |  | 1 [1; 1] |  |
| Staphylinidae | 1,5 [1; 2,5] |  | 2 [2; 2] |  |
| Curculionidae | 1 [1; 1] |  | 0 |  |
| Other Coleoptera | 1 [1; 1] |  | 0 |  |
| Diptera (larvae) | 2 [1; 17,25] |  | 0 |  |
| Aphididae | 0 |  | 1 [1; 1] |  |
| Cicadellidae | 0 |  | 1 [1; 1] |  |

A high degree of heterogeneity in the spatial distribution of the population of soil invertebrates was observed (Table 1). The total number of invertebrates encountered in the samples ranged from 1 to 22 specimens in 2019 and from 0 to 6 in 2020, with an interquartile range of 6.75 in 2019 and 3 in 2020. The distribution of the number of arthropod specimens per sample differed from normal. Geobionts were predominantly represented by larvae and adults of Coleoptera families Carabidae, Staphylinidae, Elateridae, and Scarabaeidae, as well as by Diptera larvae. Among herbivores, Coleoptera (Curculionidae) and Hemiptera (Aphididae, Miridae, Cicadellidae) were prevalent. In June, at all plots, regardless of the year, specialized phytophages of legumes of the genus *Sitona* Germar, 1817 (Curculionidae) and ground beetles (Carabidae) were prevalent. The latter were mainly represented by two species, *Bembidion quadrimaculatum* (Linnaeus, 1761) and *B. properans* (Stephens, 1828), which are usual inhabitants of arable land in agrocenoses (Kryzhanovskii, 1983).

During the experiments in both 2019 and 2020, a total of 29 groups of soil invertebrates were identified in the samples. Among soil mesofauna, Coleoptera imagoes and larvae (Carabidae, Staphylinidae, Scarabaeidae, Elateridae) and Diptera larvae were prevalent. Coleoptera (Curculionidae) and Hemiptera (Aphididae, Miridae, Cicadellidae) were prevalent among herbage inhabitants (Table 2). Representatives of other invertebrate taxa occurred sporadically in the samples and, cumulatively for all of the experimental sites, constituted from 1% to 16% of the total number of invertebrates in each count. The number of the soil mesofauna invertebrates and herbivores identified in the experimental plots during the study, in most cases, did not depend on the type of treatment. A decrease in the total number of specimens in the plots treated with *M. robertsii* was noted only in August 2020 at the level of marginal significance (p = 0.06, Table 3). Using the examples of the most numerous taxa found in 2019 (June/August – 7 out of 8 groups), and 2020 (June/August – 7 out of 9 groups), we show that, in the overwhelming majority of cases, the treatment with *M. robertsii* did not have a significant effect on the abundance of soil invertebrates (Fig. 2; Table 3).

**Table 2.** Relative proportions of invertebrates of different taxonomic and functional groups at the plots with seeds treated with the entomopathogenic fungus *Metarhizium robertsii* (Mr) and in the control plots (C). The order of the groups corresponds to the total abundance of the group for the entire duration of the experiment.

| N | Taxa | Month | 2019 |  | 2020 |  |
| --- | --- | --- | --- | --- | --- | --- |
|  |  |  | C | Mr | C | Mr |
| 1 | Aphididae | June | <0,01 | <0,01 | 0,03 | 0,11 |
|  |  | August | 0,43 | 0,44 | 0,28 | 0,35 |
| 2 | Curculionidae | June | 0,39 | 0,42 | 0,24 | 0,20 |
|  |  | August | 0,04 | 0,05 | 0,02 | <0,01 |
| 3 | Carabidae | June | 0,26 | 0,25 | 0,15 | 0,08 |
|  |  | August | 0,06 | 0,03 | 0,13 | 0,06 |
| 4 | Heteroptera | June | 0,01 | 0,01 | 0,01 | 0,03 |
|  |  | August | 0,16 | 0,10 | 0,12 | 0,10 |
| 5 | Araneae | June | 0,04 | 0,01 | 0,02 | 0,05 |
|  |  | August | 0,03 | 0,01 | 0,11 | 0,16 |
| 6 | Carabidae<br>(larvae) | June | 0,03 | 0,05 | 0,02 | 0,03 |
|  |  | August | 0,02 | 0,09 | 0,03 | 0,03 |
| 7 | Other Coleoptera | June | 0,02 | 0,02 | 0,03 | 0,02 |
|  |  | August | 0,08 | 0,06 | 0,05 | 0,02 |
| 8 | Cicadellidae | June | 0 | 0 | 0 | 0,01 |
|  |  | August | 0,02 | 0,03 | 0,08 | 0,07 |
| 9 | Staphylinidae | June | 0,07 | 0,02 | 0,02 | 0,01 |
|  |  | August | 0,04 | 0,05 | 0,04 | <0,01 |
| 10 | Scarabaeidae<br>(larvae) | June | 0,01 | 0,01 | 0 | 0 |
|  |  | August | 0,07 | 0,07 | 0,02 | 0,05 |
| 11 | Elateridae<br>(larvae) | June | 0,11 | 0,10 | 0,02 | 0,02 |
|  |  | August | 0 | 0,02 | 0 | 0 |
| 12 | Diptera<br>(larvae) | June | 0,02 | 0,11 | 0,03 | 0,01 |
|  |  | August | 0 | <0,01 | <0,01 | 0 |
| 13 | Hymenoptera | June | 0 | 0 | <0,01 | 0,01 |
|  |  | August | 0 | 0 | 0,01 | 0,02 |
| 14 | Coleoptera<br>(larvae) | June | 0 | 0 | 0 | 0 |
|  |  | August | 0 | 0 | 0,01 | 0,03 |
| 15 | Coccinellidae<br>(larvae) | June | 0 | 0 | 0 | 0 |
|  |  | August | 0,01 | <0,01 | 0,02 | 0,01 |
| 16 | Coccinellidae | June | 0,01 | 0,01 | 0 | 0 |
|  |  | August | 0 | <0,01 | 0,02 | 0,01 |
| 17 | Enchytraeidae | June | 0 | 0 | <0,01 | 0,02 |
|  |  | August | 0 | <0,01 | 0,01 | 0 |
| 18 | Anthicidae | June | 0 | 0 | 0 | 0 |
|  |  | August | 0 | 0 | 0,01 | 0,02 |
| 19 | Lepidoptera<br>(larvae) | June | 0 | 0 | 0,01 | 0,01 |
|  |  | August | 0,01 | 0,01 | 0 | <0,01 |
| 20 | Curculionidae<br>(larvae) | June | 0 | 0 | 0 | <0,01 |
|  |  | August | 0 | <0,01 | <0,01 | 0,03 |
| 21 | Neuroptera<br>(larvae) | June | 0 | 0 | 0 | 0 |
|  |  | August | 0,02 | 0,01 | 0,01 | 0 |
| 22 | Geophilina | June | 0,01 | <0,01 | 0 | 0,01 |
|  |  | August | 0,01 | 0 | 0 | 0,01 |
| 23 | Formicidae | June | 0 | 0 | 0 | 0 |
|  |  | August | 0,01 | <0,01 | 0,02 | 0 |
| 24 | Elateridae | June | 0,01 | 0,01 | 0 | 0,01 |
|  |  | August | 0,01 | 0 | 0 | 0 |
| 25 | Trombidiformes | June | 0 | 0 | 0 | <0,01 |
|  |  | August | 0,01 | 0 | <0,01 | 0,01 |
| 26 | Diptera | June | 0 | 0 | <0,01 | 0,01 |
|  |  | August | 0 | 0 | <0,01 | 0 |
| 27 | Neuroptera | June | 0 | 0 | 0 | 0 |
|  |  | August | 0 | 0 | <0,01 | 0,01 |
| 28 | Histeridae | June | 0,01 | <0,01 | 0 | 0 |
|  |  | August | 0 | 0 | 0 | 0 |
| 29 | Other Acari | June | 0 | 0 | 0 | 0 |
|  |  | August | 0 | 0 | <0,01 | 0 |
|  | **Total number of groups** | June | **15** | **15** | **14** | **19** |
|  |  | August | **17** | **19** | **24** | **20** |

**Table 3.** The abundance of invertebrates of different groups (median (25%; 75%)) at the plots with seed treatment with entomopathogenic fungi *Metarhizium robertsii* (Mr) and at the control plots (C) in June and August 2019 and 2020. Significant differences are highlighted in bold.

| Taxa | Month | 2019 |  | U | P | 2020 |  | U | P |
| --- | --- | --- | --- | --- | --- | --- | --- | --- | --- |
|  |  | C | Mr |  |  | C | Mr |  |  |
| Curculionidae | June | 12,5 [9,5; 21,5] | 19,0 [12,5; 23,3] | 5,0 | 0,471 | 6,0 [4,0; 11,0] | 8,0 [5,0; 8,5] | 10,5 | 0,754 |
|  | August | 2,0 [1,3; 2,8] | 3,0 [0,8; 3,8] | 7,0 | 0,885 | - | - | - | - |
| Carabidae | June | 10,5 [7,0; 11,8] | 12,0 [5,75; 14,5] | 5,5 | 0,564 | 3,0 [0,5; 9,0] | 3,0 [1,5; 4,0] | 12,0 | 1,00 |
|  | August | 1,0 [0,3; 7,8] | 1,0 [0,3; 4,0] | 7,5 | 1,00 | 9,0 [2,0; 14,5] | 3,0 [1,5; 4,0] | 6,0 | 0,210 |
| Elateridae (larvae) | June | 4,5 [1,8; 5,8] | 4,5 [0,8; 7,5] | 7,5 | 1,00 | - | - | - | - |
|  | August | - | - | - | - | - | - | - | - |
| Diptera (larvae) | June | 1,0 [0,3; 1,0] | 4,0 [2,5; 7,8] | **0,0** | **0,030** | 1,0 [0; 1,5] | 0 [0; 0,5] | 5,00 | 0,144 |
|  | August | - | - | - | - | - | - | - | - |
| Carabidae (larvae) | June | 1,0 [0,3; 2,5] | 2,0 [1,3; 2,8] | 4,5 | 0,387 | 1,0 [0; 1,0] | 1,0 [0; 2,0] | 9,5 | 0,602 |
|  | August | 0,5 [0; 1,8] | 2,0 [0,5; 12,5] | 4,0 | 0,312 | 1,0 [0; 4,5] | 1,0 [0; 2,5] | 9,0 | 0,531 |
| Staphylinidae | June | 2,0 [1,3; 4,3] | 1,0 [0; 2,0] | 4,0 | 0,312 | - | - | - | - |
|  | August | 1,5 [1,0; 2,8] | 3,0 [0,8; 4,5] | 5,0 | 0,470 | 3,0 [1,5; 4,0] | 0 [0; 0,5] | 3,0 | 0,060 |
| Araneae | June | 1,5 [0,3; 2,8] | 0 [0; 0,8] | 3,0 | 0,194 | 0 [0; 1,5] | 2,0 [1,0; 2,0] | 5,5 | 0,175 |
|  | August | - | - | - | - | 7,0 [4,0; 9,5] | 7,0 [5,0; 8,5] | 11,5 | 0,917 |
| Aphididae | June | - | - | - | - | 0 [0; 2,0] | 3,0 [2,0; 6,0] | **2,0** | **0,037** |
|  | August | 18,0 [11,0; 34,0] | 25,5 [8,8; 37,8] | 7,0 | 0,875 | 15,0 [8,0; 30,0] | 16,0 [12,0; 18,5] | 12,0 | 1,00 |
| Heteroptera | June | - | - | - | - | 0 [0; 1,0] | 1,0 [0,5; 2,0] | 5,5 | 0,15 |
|  | August | 4,5 [4,0; 15,5] | 5,0 [3,5; 7,3] | 8,0 | 0,885 | 6,0 [5,5; 10,5] | 4,0 [1,0; 8,0] | 6,0 | 0,210 |
| Scarabaeidae (larvae) | June | - | - | - | - | - | - | - | - |
|  | August | 0,5 [0; 10,0] | 0,5 [0; 11,5] | 5,0 | 0,470 | 0 [0; 3,0] | 0 [0; 5,0] | 11,5 | 0,91 |
| Cicadellidae | June | - | - | - | - | - | - | - | - |
|  | August | 0,5 [0; 2,5] | 2 [0,5; 2,8] | 5,5 | 0,564 | 4,0 [2,0; 8,5] | 3,0 [2,0; 4,0] | 8,5 | 0,465 |
| Coleoptera (larvae) | June | - | - | - | - | - | - | - | - |
|  | August | - | - | - | - | 1,0 [0; 1,0] | 0 [0; 3,0] | 12,0 | 1,00 |
| **The number of specimens** | June | 33,5 [31,3; 47,0] | 41,0 [31,3; 59,0] | 56,5 | 0,387 | 15,0 [10,0; 25,5] | 17,0 [16,0; 28,5] | 93,0 | 0,431 |
|  | August | 49,0 [41,5; 56,5] | 55,5 [29,0; 79,0] | 71,0 | 0,977 | 64,0 [49,5; 78,5] | 48,0 [33,5; 52,0] | **63,5** | **0,044** |
| **The number of groups** | June | 9,5 [6,8; 10,8] | 7,5 [7,0; 8,0] | 3,5 | 0,24 | 6,0 [5,0; 8,0] | 8,0 [6,5; 10,0] | 5,0 | 0,14 |
|  | August | 10,5 [10,0; 11,8] | 11,5 [8,0; 12,8] | 4,5 | 0,37 | 14,0 [12,5; 15,5] | 11,0 [8,5; 15,0] | 7,0 | 0,29 |

**Fig. 2.**
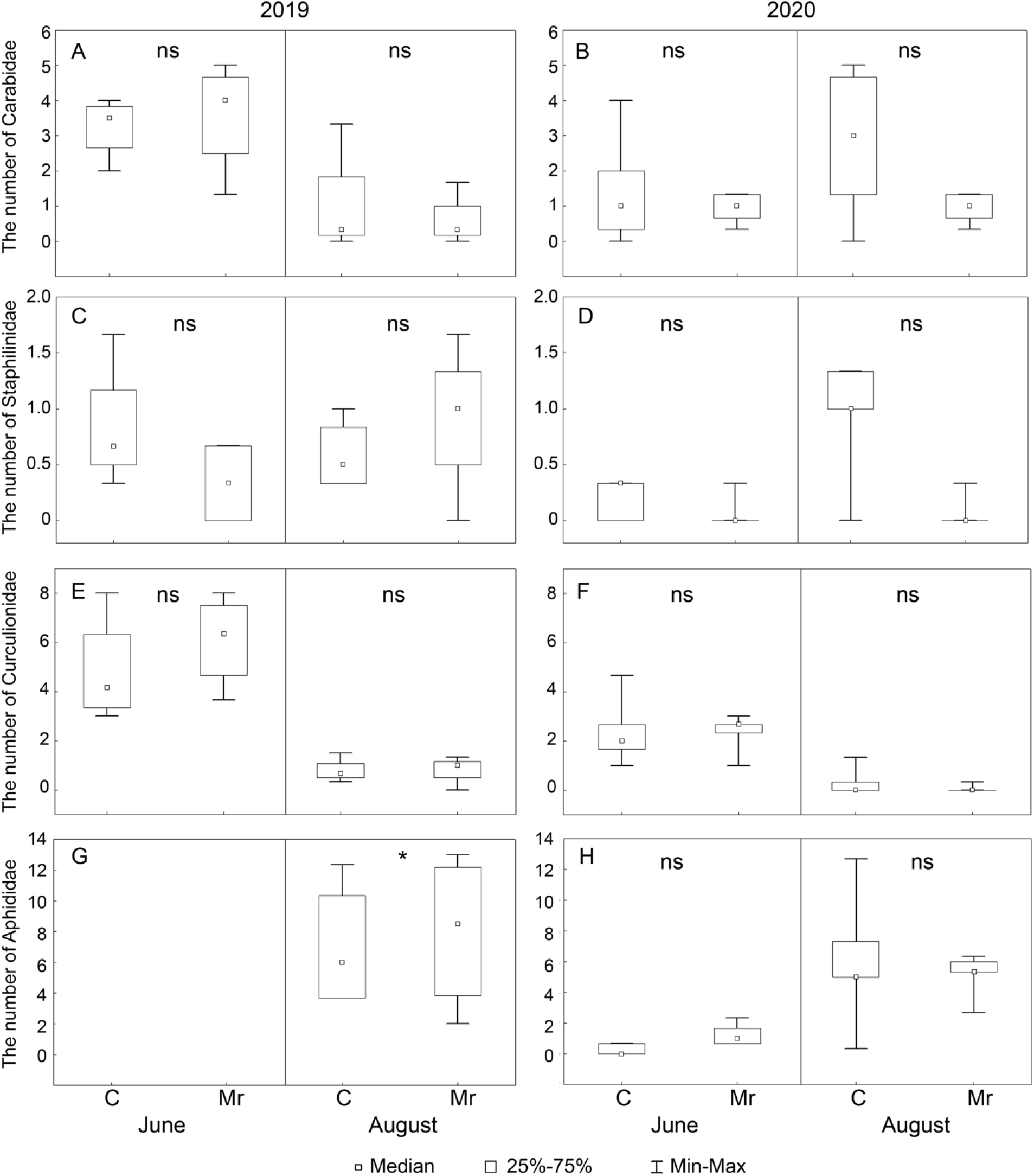
Abundance of representatives of key groups of the soil mesofauna and herbivores (specimens / sample) at the plots with seed treatment with the entomopathogenic fungus *Metarhizium robertsii* (Mr) and at the control plots (C) in June and August 2019 and 2020. Taxonomic groups: A, B - Carabidae; C, D - Staphilinidae; E, F - Curculionidae; G, H - Aphididae. Mann-Whitney test: * p <0.05; ns - p> 0.05. ns-significance

Significant positive effects of treatment were noted for two groups. In June 2019, the abundance of fly (Diptera) larvae was significantly higher in the plots with seed treatment with *M. robertsii* compared to the control (Table 3). In June 2020, a significant increase in the number of aphids in samples collected from the plots with treated seed was noted (Fig. 2; Table 3).

#### Population diversity

Soil fauna communities at plots with different treatment types included 14 to 24 invertebrate groups (Table 2), with no significant effect of treatment on the number of taxa in both 2019 and 2020 (Table 3). In most cases (with the notable exception of June 2020), treatment had no significant effect on the species diversity and species evenness in the invertebrate communities at the experimental sites (Table 4).

**Table 4.** Diversity indices in invertebrate communities at the experimental plots with seed treatment with entomopathogenic fungus *Metarhizium robertsii* (Mr) and at the control plots (C) in different years. Different letters in the lines indicate significant differences for individual years (permutation criterion: p <0.05).

| Indices | Month | 2019 |  | 2020 |  |
| --- | --- | --- | --- | --- | --- |
|  |  | C | Mr | C | Mr |
| Shannon index | June | 1,791<sup>a</sup> | 1,682<sup>a</sup> | **1,801<sup>a</sup>** | **2,164<sup>b</sup>** |
|  | August | 1,964<sup>a</sup> | 2,045<sup>a</sup> | 2,421<sup>a</sup> | 2,268<sup>a</sup> |
| Simpson index | June | 0,759<sup>a</sup> | 0,741<sup>a</sup> | **0,747<sup>a</sup>** | **0,827<sup>b</sup>** |
|  | August | 0,769<sup>a</sup> | 0,772<sup>a</sup> | 0,863<sup>a</sup> | 0,827<sup>a</sup> |
| Buzas-Gibson Evenness | June | 0,461<sup>a</sup> | 0,448<sup>a</sup> | 0,505<sup>a</sup> | 0,484<sup>a</sup> |
|  | August | 0,413<sup>a</sup> | 0,407<sup>a</sup> | 0,450<sup>a</sup> | 0,460<sup>a</sup> |
| Berger-Parker | June | 0,389<sup>a</sup> | 0,417<sup>a</sup> | 0,419<sup>a</sup> | 0,330<sup>a</sup> |
|  | August | 0,429<sup>a</sup> | 0,440<sup>a</sup> | 0,284<sup>a</sup> | 0,352<sup>a</sup> |

At the end of June 2020, the species diversity and abundance of invertebrates were significantly higher at the plots with *M. robertsii* treatment: the values of the Shannon and Simpson indices on treated plots were significantly higher than in the control.

The distributions of variants of the invertebrate population along the ordination plane in the experimental and control plots were similar in June, August 2019, and June 2020, forming symmetrical clouds around the origin of coordinates (Fig. 3. A, B, C). In August 2020, the distribution of points formed by the experimental sites was much denser, even though the centers of the distributions in the experiment and control were the same (Fig. 3, D). This may indicate greater uniformity in the population structure at the experimental sites.

**Fig. 3.**
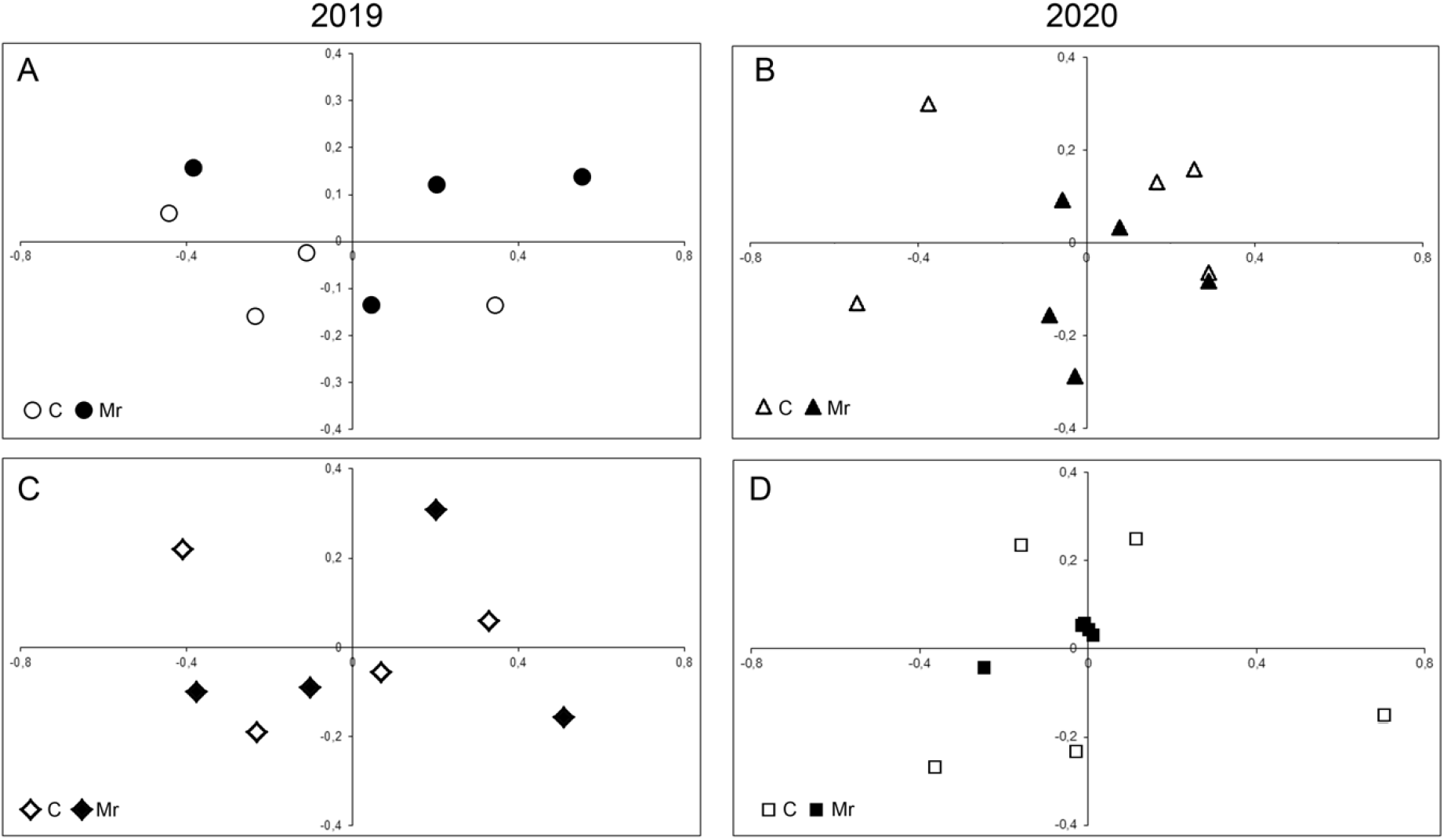
Ordination (Euclidean distance, multidimensional scaling) of variants of the invertebrate populations at the experimental plots in June 2019 (A), June 2020 (B), August 2019 (C) and August 2020 (D). Control plots are marked with white icons, while the treated plots with *M. robertsii* are marked with black icons.

### Aphids

Aphids (Hemiptera: Aphididae) of four species from three genera were detected on the beans: *Acyrthosiphon pisum* (Harris, 1776), *Megoura viciae* Buckton, 1876, *Aphis fabae* Scopoli, 1763 and *Aphis spiraecola* Patch, 1914. *A. pisum* were prevalent both in 2019 and 2020 (Table 5).

**Table 5.**
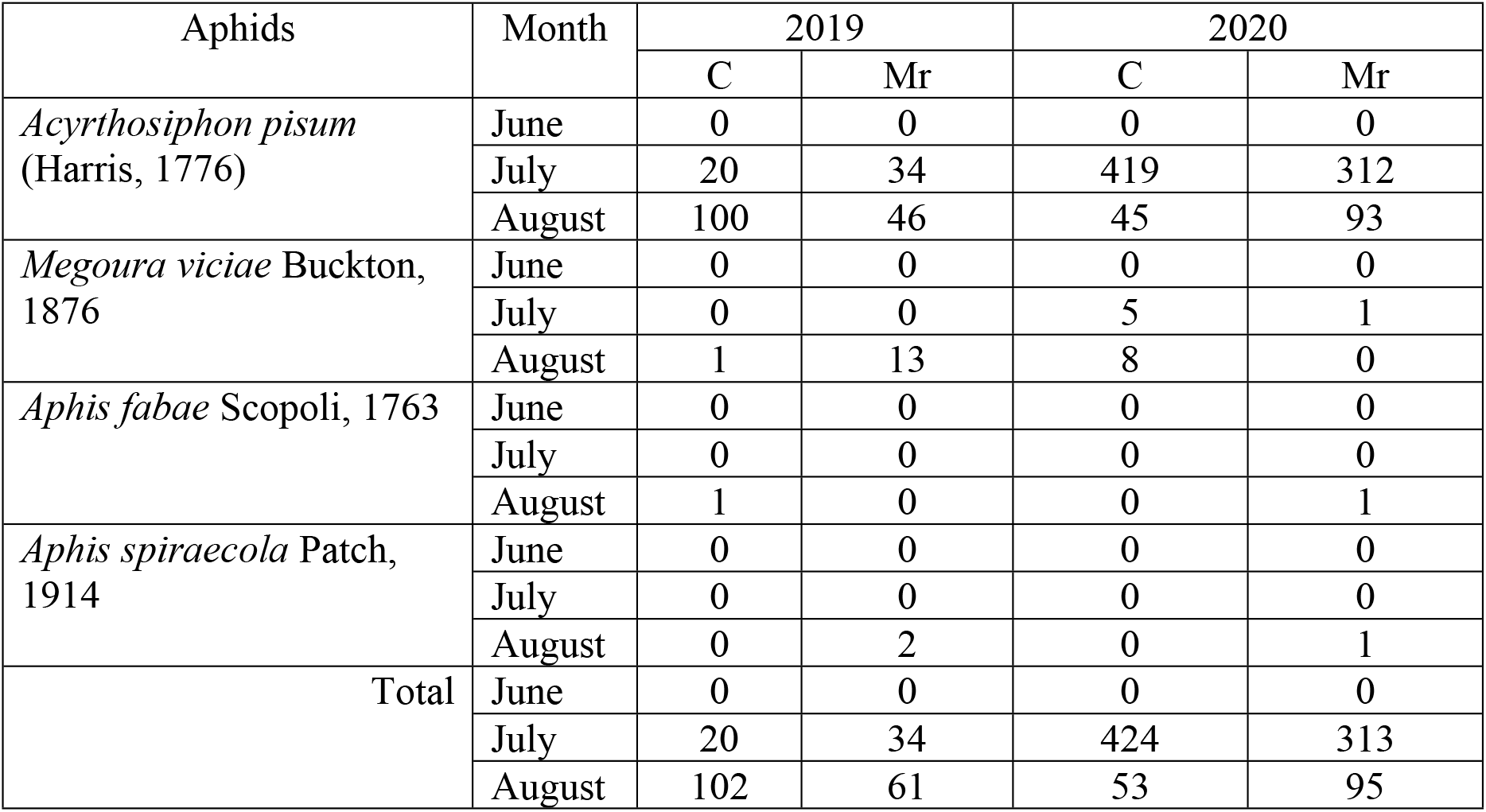
The number of aphids collected at the experimental sites with seed treatment with the entomopathogenic fungus *Metarhizium robertsii* (Mr) and at the control sites (C) in different years.

Colonies of aphids *Megoura viciae* were rare and few. Single aphids *Aphis fabae* and *A. spiraecola* were observed at the experimental plots only in August.

In the first ten days of June (plant branching phase), aphids were not found on the examined plants in 2019 or in 2020. In July (budding phase), aphids of two species (*A. pisum, M. viciae*), with absolute prevalence of *A. pisum*, were detected at the experimental plots (Table 5). The ratio of *A. pisum* to the total number of aphids collected during this period was 100% in 2019 and 99.19% in 2020 (Table 5). There were no significant differences between the treated and the control plots in terms of the proportion of plants infested with aphids (Mann-Whitney test: 2019, U = 7.00, p = 0.88; 2020, U = 7.50, p = 0.35; Fig. 4) and in terms of the overall number of aphids on individual plants (Mann-Whitney criterion: 2019: U = 5.00, p = 0.47; 2020: U = 9.00, p = 0.53; Fig. 5).

**Fig. 4.**
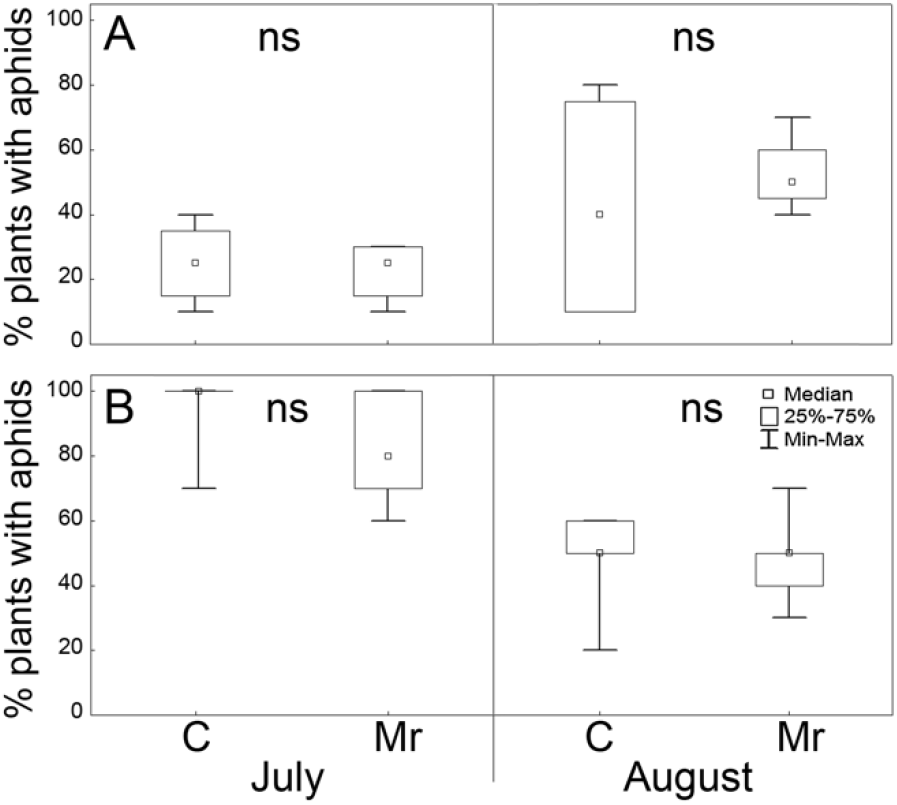
Proportion of *Vicia faba* L. plants with aphid colonies from plots with beans treated with the entomopathogenic fungus *Metarhizium robertsii* (Mr) and from the control plots (C) in July and August 2019 (A) and 2020 (B). Mann-Whitney test: ns - p> 0.05.

**Fig. 5.**
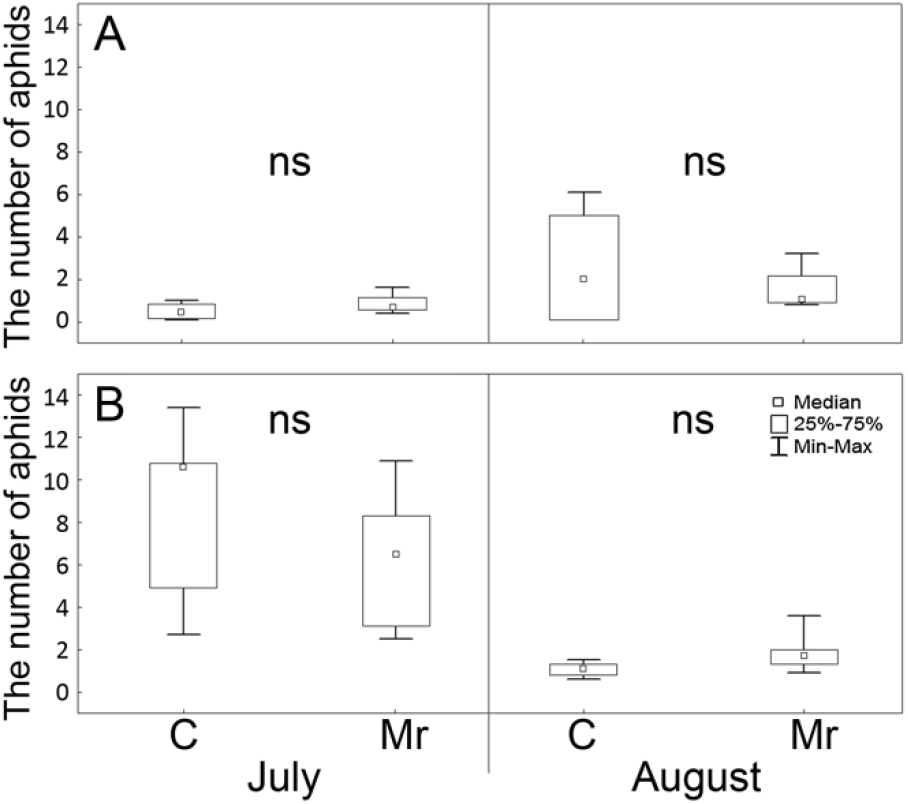
The number of aphids on the *Vicia faba* L. plants with the beans during seed treatment with the entomopathogenic fungus *Metarhizium robertsii* (Mr) and the control (C) in July and August 2019 (A) and 2020 (B). Mann-Whitney test: ns – no significance, * – p> 0.05.

A similar situation was observed at the end of the season. In August (seed maturation phase), aphids of 4 species with the prevalence of *A. pisum* were observed at the experimental plots (Table 5). The ratio of *A. pisum* to the total number of aphids collected during this period in 2019 and in 2020 was 89.57% and 93.24%, respectively. During this period, aphids of 2 to 3 species were encountered at the plots with different types of treatment. There were also no significant differences between the plots treated with *M. robertsii* and the control plots, both in the proportion of plants infested with aphids (Mann-Whitney test: 2019: U = 7.5, p = 1; 2020: U = 11.0, p = 0.83; Fig. 4), and in terms of the number of aphids on plants (Mann-Whitney criterion: 2019: U = 796, p = 0.969; 2020: U = 4.5, p = 0.12; Fig. 5).

In summary, according to the results of this two-year experiment, treatment of seeds of *Vicia faba* L. beans with the entomopathogenic fungus *M. robertsii* did not significantly alter the proportion of plants with aphids or the density of aphid colonies on individual plants.

### Leaf miner flies

At all stages of the experiment in 2019 and 2020 the leaves of the beans *Vicia faba* L. were damaged by the leaf miner fly *Liriomyza bryoniae* (Kaltenbach, 1858) (Agromyzidae). According to the data on the imago hatching from pupae, the rate of infection of larvae with parasitoids at the experimental plots was quite high both in 2019 and in 2020, namely, 64.7 % and 62.5 %, respectively.

Treatment of bean seeds with the entomopathogenic fungus *M. robertsii* did not significantly affect the rate of infestation of plant leaves with miners in most cases, with the exception of August 2020 (Fig. 6). During this period, the proportion of infected leaves in treated plants was significantly higher than in control (Mann-Whitney test: U = 0.00, p = 0.009).

The degree of leaf blades damaged by the miner at the experimental plots (and, in general, at all plots, regardless of the treatment) in different years was quite low and ranged from 1 to 3 mines per leaf.

The percentage of infected leaves in the middle part of the plant did not differ significantly in different years (Mann-Whitney test: 2019: 8.83 [7.24; 14.24]; 2020:

13.21 [10.26; 15.16]; U = 27.0, p = 0.27). The percentage of leaf infestation in the upper part of the plant was significantly higher in August 2019 compared to 2020 (Mann-Whitney test: 2019: 30.63 [29.05; 40.55]; 2020: 8.24 [7.54; 15.31]; U = 0.0, p = 0.0004). Due to the registration of infestation of the lower leaves in 2019 in July instead of June, a comparative analysis for the lower part in this case was not carried out.

## Discussion

The presence of common soil fungi, such as entomopathogenic ascomycetes *Metarhizium robertsii*, in the soil in the case of agricultural cultivation may lead to desirable effects on plants. However, the colonization of plants by the fungi can affect the composition and structure of invertebrate communities. The effect of entomopathogenic fungi on non-target arthropods has not been sufficiently studied, with studies being limited to laboratory conditions or direct-contact conditions of the invertebrates with fungal propagules. This work aimed to close this gap in knowledge by analyzing the effect of colonization of broad been seeds on invertebrate communities in the field conditions, as would be used in agriculture.

Our two-year analysis showed that inoculation of beans with the conidia of entomopathogenic fungus *M. robertsii* does not lead to significant alterations in the invertebrate community. Nevertheless, we detected some short-term significant changes in the abundance of species and the number of phytophages and saprophages, as well as an increased abundance of arthropods. The composition of soil invertebrates found at the experimental site was typical for agricultural fields of chernozems (36).

The treatment of broad beans with the *M. robertsii* did not significantly affect the composition or the abundance of the soil dwellers and herbivores in most cases. Most invertebrates rarely have contact with the soil (grass and soil surface inhabitants) and therefore did not have contact with fungal propagules. Many of them have a very low abundance which is insufficient for the entomopathogenic effect to be revealed by the assay. However, this assumption requires careful verification. At the same time, in June 2019, a positive effect of fungal treatment on the abundance of soil dipteran saprophage larvae was noted. This may be due to the tendency of fly larvae to form aggregations in places with an abundance of food, one of which, due to random reasons, existed on plots with *M. robertsii*. In addition, the positive effect of the treatment was revealed at the end of June 2020 for aphids. This could be due to the accelerated growth and development of plants after seed treatment with the fungus, which makes plants more attractive to winged migrants during their dispersal in June. Similarly, the higher species abundance of arthropods at the sites treated with *M. robertsii* in June 2020 could be explained by the faster development of plants after seed treatment with the fungus (22).

### Aphids

All aphid species we found at the experimental plots (*Acyrthosiphon pisum, Megoura viciae, Aphis fabae*, and *A. spiraecola*) are widespread and common both for natural habitats and for agroecosystems (31)(2020). Aphids of both species of the genus *Aphis* are found on a wide range of secondary host plants, including many agricultural plants (29). *Acyrthosiphon pisum* and *Megoura viciae* prefer plants from the Fabaceae family, while *A. pisum* belongs to dangerous pests of agricultural crops. *A. pisum* was prevalent at the experimental plots at all stages of the experiment.

The absence of aphids on the surveyed plants at the first ten days of June 2019 and 2020 could be explained by regional specifics. During this period, winged migrants which did not yet manage to colonize experimental plants dispersed. At the experimental sites during this period, there were only a few winged specimens. Two weeks later, at the end of June, a low abundance of aphids was observed, and a positive effect of seed treatment on the number of aphids during the dispersal of winged migrants was revealed. However, later these differences disappeared later.

In July and August, aphids were common on bean plants. The number of aphids in individual colonies reached up to 70 specimens both on treated and control plots. It is known that seed treatment and subsequent colonization of plants with entomopathogenic fungi can lead to a significant decrease in density of aphids on plants (37). However, according to our two-year field experiment, the treatment of *Vicia faba* beans with *M. robertsii* did not show a significant effect on the proportion of plants with aphids and on the density of aphid colonies on individual plants both in the middle (plant budding phase) and end of the field season (bean ripening phase). This could be due to the low level of colonization of internal plant tissues by the fungus. Earlier it was shown that when treating cotton seeds with fungi *Purpureocillium lilacinum* and *Beauveria bassiana*, with the level of colonization of about 50%, significant differences were established only in a comparative analysis of colonized and non-colonized plants, while when analyzing the general effect of treatment, significant differences could not be detected (37). In our case, we could not assess the effect of colonization due to the low proportion of plants inhabited by the fungus (22).

### Leaf miner flies

The tomato leafminer *Liriomyza bryoniae* (Kaltenbach, 1858) is included in the list of quarantine species of the International Plant Protection Convention (26). *L. bryoniae* is a polyphage (38). Our data indicate a low level of infection of plants with *L. bryoniae* larvae at the experimental site, and, most likely, the absence of a significant effect of this species on the yield of beans. With a fairly low degree of damage to leaf blades (1–3 mines per leaf), the proportion of infected leaves at different stages of plant development varied from 4.55 % to 87.43 % in 2019 and from 0 % to 21.64 % in 2020.

The low level of infestation of broad beans with *L. bryoniae* in this case may be due to the high (more than 60 %) level of infestation of fly larvae with hymenopteran parasitoids, which can have a substantial impact on the population of *L. bryoniae* in natural conditions (39). It is known that entomopathogenic fungi can lead to significant negative effect on the population of mining flies (40,41). Under the conditions of our experiment, the treatment of beans with *M. robertsii* fungi did not cause changes in the degree of infestation of plant leaves. Moreover, in August 2020, the infestation of the upper leaves upon treatment with *M. robertsii* was significantly higher compared to control. This probably occurred due to the low level of plant colonization with the fungus under natural conditions. Laboratory experiments have shown that mortality rate of flies depends both on the isolate and the degree of plant colonization by the fungus (18,19). It is also possible that plants with higher plant biomass were more attractive to miners.

## Conclusions

1. Treatment of *Vicia faba* beans with the entomopathogenic fungus *M. robertsii* in most cases did not cause significant changes neither on the total abundance of soil dwellers and grass stand inhabitants, nor on the abundance of the most abundant taxa (Coleoptera: Carabidae, Staphylinidae, Elateridae, Scarabaeidae, Curculionidae; Diptera larvae; Hemiptera: Miridae, Cicadellidae, Aphididae). The effects revealed for certain groups were slight. A positive effect of treatment on population density was registered only for soil saprophagous flies larvae in June 2019.
2. Among aphids, *Acyrthosiphon pisum, Megoura viciae, Aphis fabae*, and *Aphis spiraecola* were registered with a predominance of *A. pisum*. Seed treatment with the entomopathogenic fungus *M. robertsii* did not significantly affect the proportion of plants inhabited by aphids or the density of aphid colonies on individual plants throughout the season.
3. At all stages of the experiment, the bean leaves were damaged by the mining flies *Liriomyza bryoniae* and high level of infestation with Hymenoptera was observed (more than 60 %). With a fairly low degree of damage to leaf blades (1-3 mines per leaf), the proportion of infected leaves varied from 4.55 % to 87.43 % in 2019 and from 0 % to 21.64 % in 2020. Bean seed treatment with *M. robertsii* did not significantly affect the degree of infestation of plant leaves by larvae of the *L. bryoniae*.

In summary, the treatment of broad bean seeds with the entomopathogenic fungus *M. robertsii* in agroecosystems of Western Siberia did not significantly affect the non-target groups of arthropods (typical for bean field) or the main pests of beans (aphids and miner flies).

